# Grandmaternal allergen exposure causes distinct epigenetic trajectories in offspring associated with airway hyperreactivity and inflammation

**DOI:** 10.1101/2022.01.25.477760

**Authors:** Katie M. Lebold, Madeline Cook, Alexandra B. Pincus, Kimberly A. Nevonen, Brett A. Davis, Lucia Carbone, Gina N. Calco, Allison D. Fryer, David B. Jacoby, Matthew G. Drake

## Abstract

Maternal asthma increases childhood asthma risk through multiple mechanisms including epigenetic regulation of asthma-associated genes. DNA methylation is one form of epigenetic regulation that is both inherited and modified by environmental exposures throughout life. In this study, we tested whether grandmaternal house dust mite (HDM) allergen exposure altered airway physiology and inflammation, as well as DNA methylation in both airway epithelium and airway sensory neurons of second-generation offspring. Grandmaternal allergen exposure induced a limited number of epigenetic changes in offspring at baseline that were not associated with increased airway reactivity or inflammation. In contrast, grandmaternal allergen exposure significantly altered offspring’s response to HDM sensitization and challenge, inducing airway hyperreactivity to inhaled serotonin, increased airway inflammation, and potentiated DNA methylation. Gene sequences susceptible to methylation after allergen sensitization, and their corresponding biological processes and enriched pathways, were unique in offspring from HDM-exposed founders, indicating that grandmaternal allergen exposure established an epigenetic trajectory in offspring at birth that directed epigenetic and physiologic responses to subsequent allergen sensitization and challenge, contributing to inheritance of asthma risk.

**SUMMARY:** Grandmaternal allergen exposure establishes an intergenerational, tissue-specific epigenetic trajectory in offspring at birth, which uniquely directs responses to allergen sensitization and challenge later in life and contributes to inheritance of asthma risk.

## INTRODUCTION

Asthma is a chronic inflammatory airway disease that often begins in childhood and is characterized by airflow obstruction and airway hyperreactivity (Pijnenburg et al., 2021). Asthma prevalence has been increasing in recent decades and is more common within families (Paaso et al., 2014). While genetic predisposition is a major contributor to asthma risk (Paller et al., 2019, Young et al., 1991), it does not fully account for rising prevalence worldwide. Recent studies indicate that maternal asthma increases childhood asthma risk more than paternal asthma (Lim et al., 2010) and better maternal asthma control during pregnancy reduces a child’s risk of asthma (Morten et al., 2018). Together, these studies suggest fetal exposure to maternal factors *in utero* may uniquely contribute to a child’s asthma risk and serve as a modifiable exposure.

Epigenetic modification, namely DNA methylation of cytosine nucleotides located within CpG sites, is one potential mechanism by which environmental exposures can alter gene expression to increase asthma risk. DNA methylation contributes to type-2 (T_H_2) T cell differentiation (Allan et al., 2012) and transcription of effector cytokines (Hosokawa et al., 2015), which correspond with increased asthma and atopy susceptibility (Barton et al., 2017, Seumois et al., 2014). Similarly, methylation changes at birth correlate with increased immunoglobulin E production throughout life (Han et al., 2021, Peng et al., 2018), possibly contributing to enhanced IgE-mediated allergen sensitization and atopy risk. The neonatal and early childhood epigenome is sensitive to maternal exposures that occur *in utero*, including maternal smoking (Joubert et al., 2016), stress (Trump et al., 2016), and asthma (Gunawardhana et al., 2014). Genome-wide and gene-specific differential methylation in early childhood predicts later development of asthma (Reese et al., 2019, den Dekker et al., 2019) and maternal asthma modifies gene-specific associations with asthma risk (DeVries et al., 2017), highlighting the impact of prenatal exposures and epigenetic changes on childhood asthma risk.

How epigenetic alternations identified at birth increase asthma risk later in life is not known. While differential genome-wide methylation is consistently found in children with asthma compared to children without asthma (Yang et al., 2017, Cardenas et al., 2019, Yang et al., 2015), differentially methylated genes are often not present at birth (Xu et al., 2018), suggesting methylation associated with asthma risk differs from that which manifest with disease onset. DNA methylation is a dynamic process that is both heritable and changes throughout life in response to environmental exposures. Allergen sensitization, a common feature in nearly two-thirds of asthmatics, correlates with broad methylation changes in atopic children (Peng et al., 2019). Similarly, exposure to allergens also leads to changes in DNA methylation in animals (Zhang et al., 2018, Peng et al., 2019, Kim et al., 2020a). Other environmental exposures associated with increased asthma risk, including air pollutants (Zhang et al., 2018) and viral infections (Spalluto et al., 2017, Pech et al., 2018), also modify DNA methylation, further confounding the relationship between epigenetic signatures at birth and signatures found in patients with established disease.

Epigenetic regulation is also tissue-specific. Most studies have utilized peripheral blood cells to compare epigenetic signatures between patients with and without asthma, however these cells may not reflect methylation changes in the lungs (Brugha et al., 2017, Lin et al., 2020). Few studies have yet to evaluate epigenetic signatures directly in the lungs in a cell-specific manner. Airway epithelial cells are of particular interest given that they line the interface between host and environment and initiate immunologic responses to inhaled allergens and other environmental exposures that contribute to both development of asthma and its exacerbations (Lambrecht and Hammad, 2012). Airway epithelium is also densely innervated by sensory neurons arising from vagal nodose-jugular ganglia, which detect inhaled stimuli and trigger reflex bronchoconstriction when activated (Drake et al., 2018, Lebold et al., 2019). Identifying epigenetic signatures in these key cells at the lung-environment interface and correlating epigenetic changes with disease features such as airway hyperreactivity and inflammation will help elucidate connections between epigenetic regulation and disease pathogenesis.

In the current study, we hypothesized that grandmaternal asthma creates an intergenerational, tissue-specific epigenetic signature that is established at birth and informs epigenetic trajectories after allergen sensitization. We tested our hypothesis by comparing CpG methylation in airway epithelial cells and vagal sensory neurons before and after HDM allergen sensitization, as well as airway physiology and inflammation after HDM sensitization and challenge, in second generation offspring descended from allergen-exposed and vehicle-exposed founding grandmothers.

## RESULTS

### Grandmaternal allergen exposure potentiates airway hyperreactivity and inflammation in second-generation offspring

Airway physiology and inflammation were assessed after HDM sensitization and challenge in second-generation offspring (F2) from HDM-exposed and vehicle-exposed grandmaternal founders (F0) (**Figure 1**). Airway reactivity to inhaled serotonin was similar between vehicle-challenged F2 offspring from F0 HDM-exposed founders (F0HDM•F2Veh) and vehicle-challenged F2 offspring from vehicle-exposed F0 founders (F0Veh•F2Veh) (**Figure 2A**), indicating that grandmaternal allergen exposure did not affect offspring’s airway reactivity at baseline. HDM sensitization and challenge of F2 mice caused airway hyperreactivity in descendants from both HDM- and vehicle-exposed F0 lineages. However, F2 airway hyperreactivity was significantly potentiated in offspring from the grandmaternal HDM-exposed lineage (F0HDM•F2HDM) compared to offspring of the vehicle-exposed F0 lineage (F0Veh•F2HDM).

**Figure 1.**
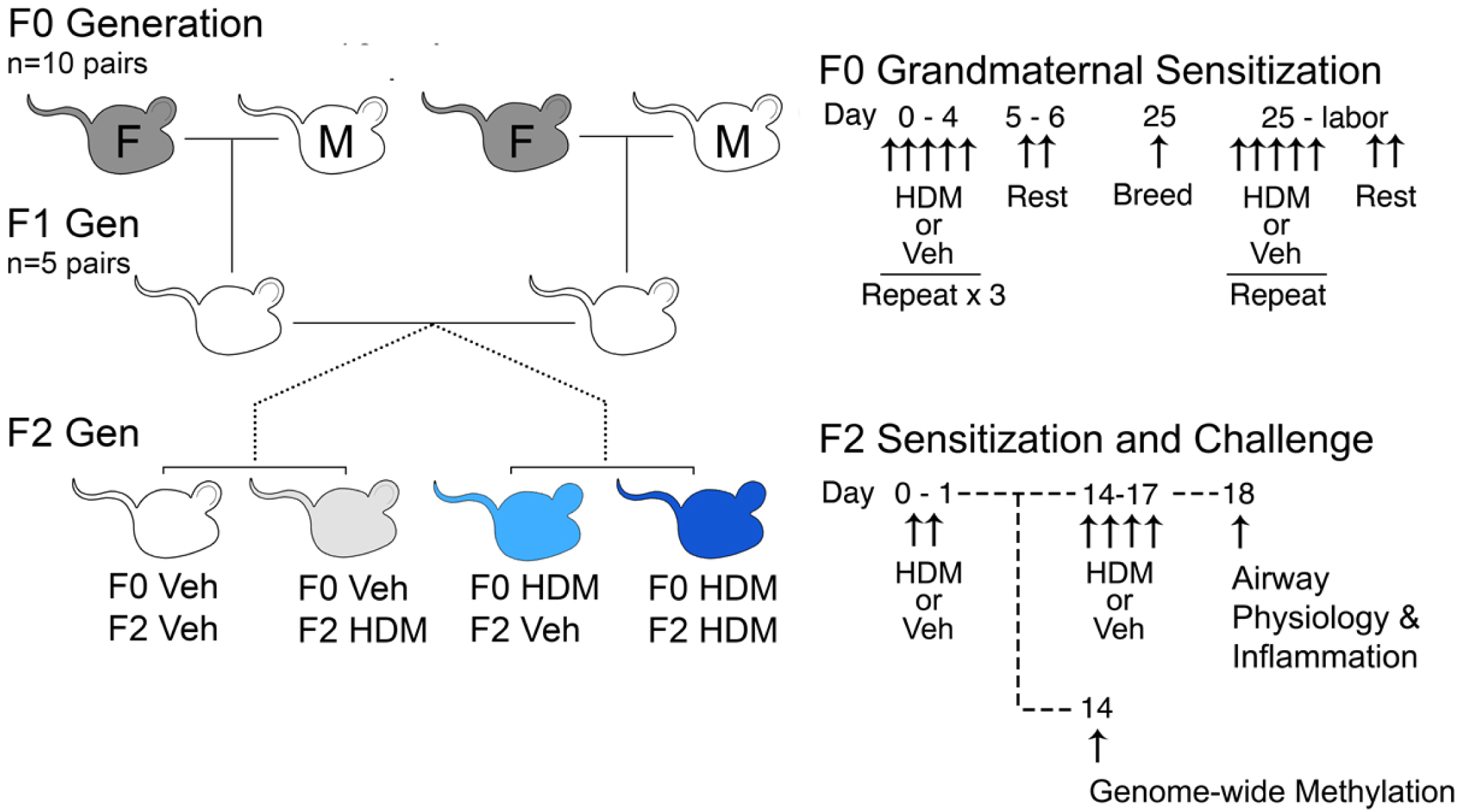
Breeding scheme and house dust mite sensitization protocol. Female F0 mice (grey mice, “F”) were exposed to intranasal house dust mite (HDM) or vehicle (Veh) for five consecutive days followed by two days of rest for 4 weeks, mated, and then exposed to HDM or vehicle 5 days per week through gestation until birth. Male F0 mice (white mouse, “M”) received no treatment. F1 male and female offspring were randomized into breeding pairs and received no treatments. Male and female F2 offspring were either sensitized to HDM or treated with Veh on days 0 and 1 and then challenged with HDM (F0Veh•F1HDM and F0 HDM•F1HDM) or vehicle (F0Veh•F1Veh and F0HDM•F1 Veh) on days 14-17. Airway physiology and bronchoalveolar lavage leukocytes were measured on protocol day 18. A subset of F2 mice were euthanized before allergen sensitization (F0Veh•F1Base and F0HDM•F1Base) and after HDM sensitization (sensitized day 0 and 1, euthanized day 14; F0Veh•F1Sens and F0HDM•F1Sens) for methylation experiments.

**Figure 2.**
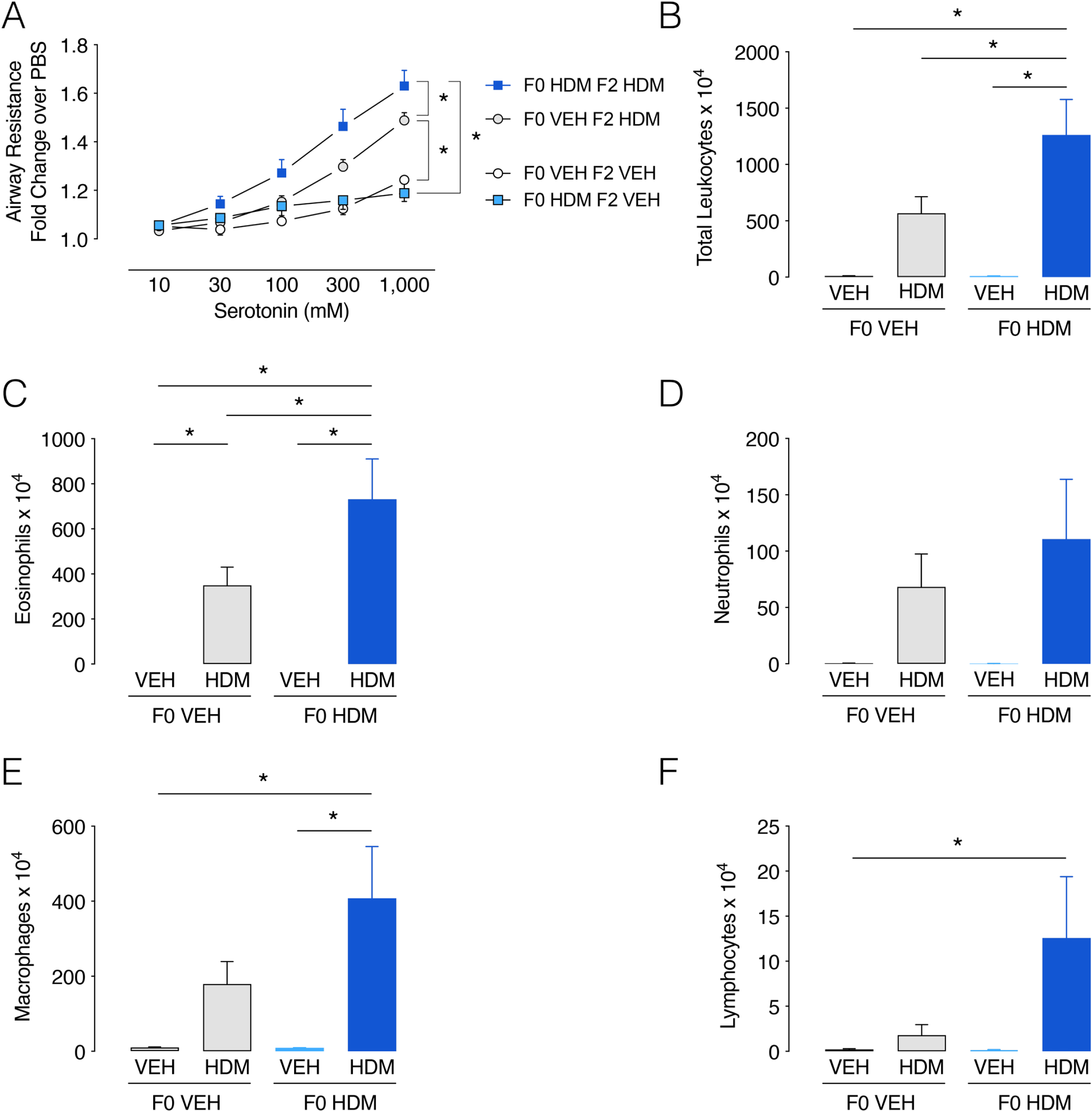
Allergen-induced airway hyperreactivity and inflammation is potentiated in second-generation offspring of HDM-exposed founders. Airway resistance was measured after escalating concentrations of aerosolized serotonin in ventilated F2 mice. Airway hyperreactivity occurred after HDM-challenge in F2 mice (F0Veh•F2HDM and F0HDM•F2HDM) compared to vehicle-challenged F2 mice (F0Veh•F2Veh and F0HDM•F2Veh). HDM-induced airway hyperreactivity was potentiated in F2 mice from HDM-exposed F0 mice (F0HDM•F2HDM) compared to F2 mice from vehicle-exposed F0 mice (F0Veh•F2HDM). Vehicle-challenged F2 offspring from both lineages had similar airway reactivity to inhaled serotonin. N = 13-20/group; *p*<0.05 based on two-way ANOVA with repeated measures. (B-F) Total airway inflammatory cells and differential cell counts were measured in bronchoalveolar lavage. Grandmaternal HDM exposure potentiated HDM-challenge induced airway inflammation in F2 offspring, including total leukocytes (B) and eosinophils (C). Neutrophils (D), macrophages (E), and lymphocytes (F) were also increased, but did not reach significance.

Similarly, airway inflammation after HDM sensitization and challenge was also augmented in the F2 generation after grandmaternal HDM exposure. Specifically, total bronchoalveolar lavage leukocytes and eosinophils were significantly increased in HDM-challenged F2 mice from HDM-exposed F0 founders (F0HDM•F2HDM) compared to HDM-challenged F2 mice from vehicle-exposed F0 founders (F0Veh•F2HDM) (**Figure 2B-F**). Macrophages, neutrophils, and lymphocytes were increased as well, but did not reach statistical significance. Grandmaternal allergen exposure did not affect second generation offspring’s airway inflammation at baseline as no differences were observed between vehicle-challenged F2 mice from the F0 HDM lineage (F0HDM•F2Veh) compared to vehicle-challenged F2 mice from the vehicle-exposed F0 lineage (F0Veh•F2Veh).

### Grandmaternal allergen exposure alters DNA methylation in allergen-naïve second generation offspring

To test whether grandmaternal allergen exposure affects offspring DNA methylation at baseline before allergen sensitization, F2 mice from HDM-exposed F0 founders (F0HDM•F2Base) were compared to F2 mice from vehicle-exposed founders (F0Veh•F2Base). In total, 290 differentially methylated regions (DMRs) were detected in airway epithelium of F0HDM•F2Base compared to F0Veh•F2Base, which overlapped 297 unique genes. Of these DMRs, 177 (61%) were hypermethylated, while 113 (39%) were hypomethylated. Most DMRs spanned intra- and intergenic DNA segments, while 18% mapped to promoters (**Figure 3**). Likewise, 1,390 differentially methylated cytosines (DMCs) were detected in airway epithelium of F0HDM•F2Base compared to F0Veh•F2Base, of which 804 (58%) were hypermethylated and 586 (42%) were hypomethylated. Again, most DMCs mapped to intra- and intergenic sequences, with 18% spanning promoters (**Figure S1**).

**Figure 3.**
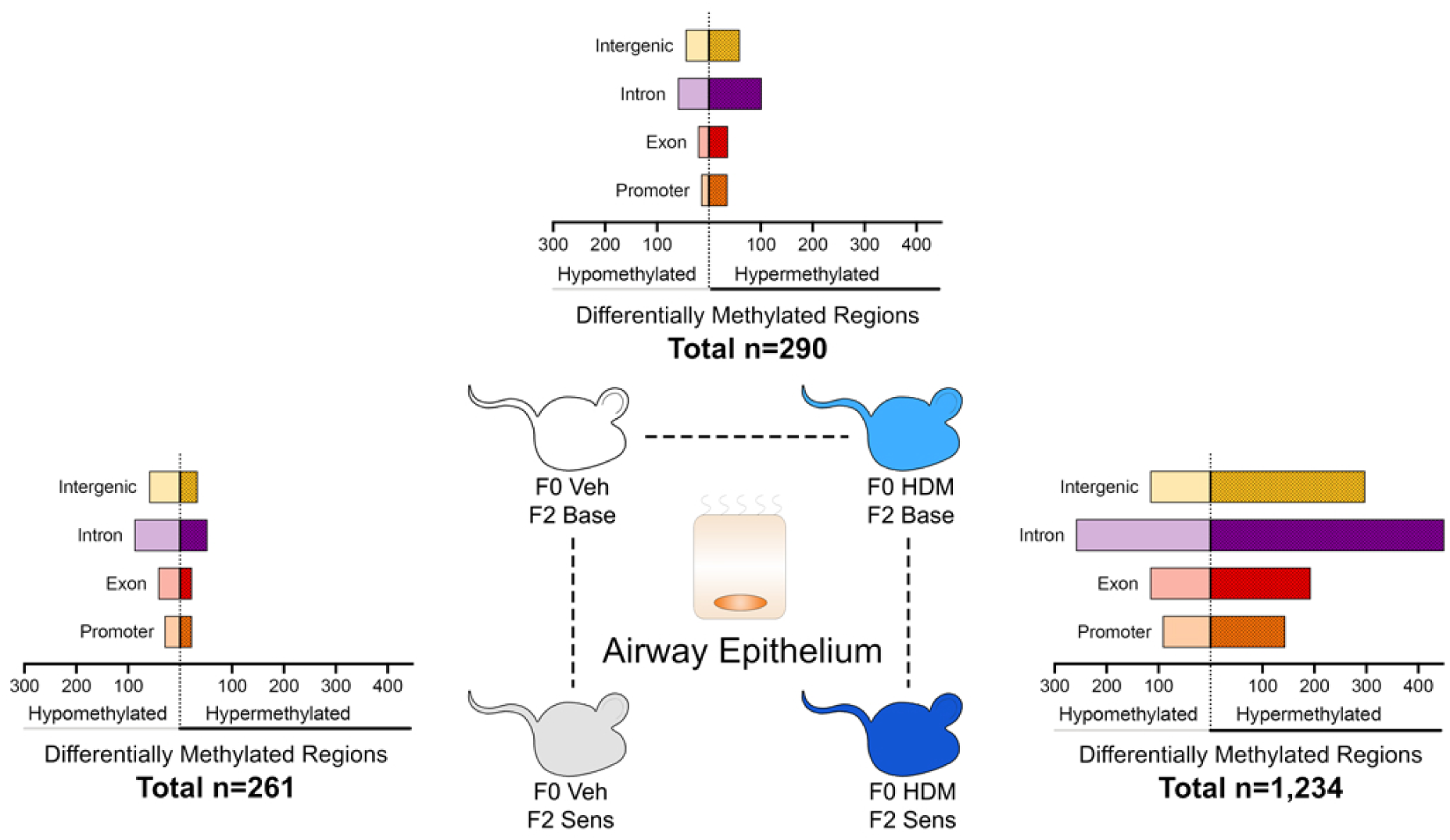
Grandmaternal allergen exposure alters airway epithelial DNA methylation at baseline and after allergen sensitization in second-generation offspring. Whole genome methylation was assessed in airway epithelium of second-generation mice (F2) before and after allergen sensitization. Grandmaternal HDM exposure alone resulted in 290 differentially methylated regions (F0 HDM•F2 Base versus F0 Veh•F2 Base) with a predominantly hypermethylated response. F2 HDM sensitization alone resulted in 261 differentially methylated regions with a predominantly hypomethylated response (F0Veh•F2Sens versus F0 Veh•F2 Base). In contrast, HDM-sensitized F2 mice from HDM-exposed F0 founders had 1,234 total differentially methylated regions with a predominantly hypermethylated response (F0 HDM•F2 Sens versus F0 HDM•F2 Base).

Gene ontology (GO) terms were used to assess for pathways and biological processes enriched in these DMRs and DMCs. For DMCs, 74 biologic processes were significantly enriched and subsequently grouped into functionally related gene clusters, including most prominently, calcium homeostasis, bone morphogenetic protein response, and cell and organ development (**Figure 4A**). No biological processes were associated with DMRs, nor were any pathways enriched within DMRs or DMCs (**Table 1**).

**Figure 4.**
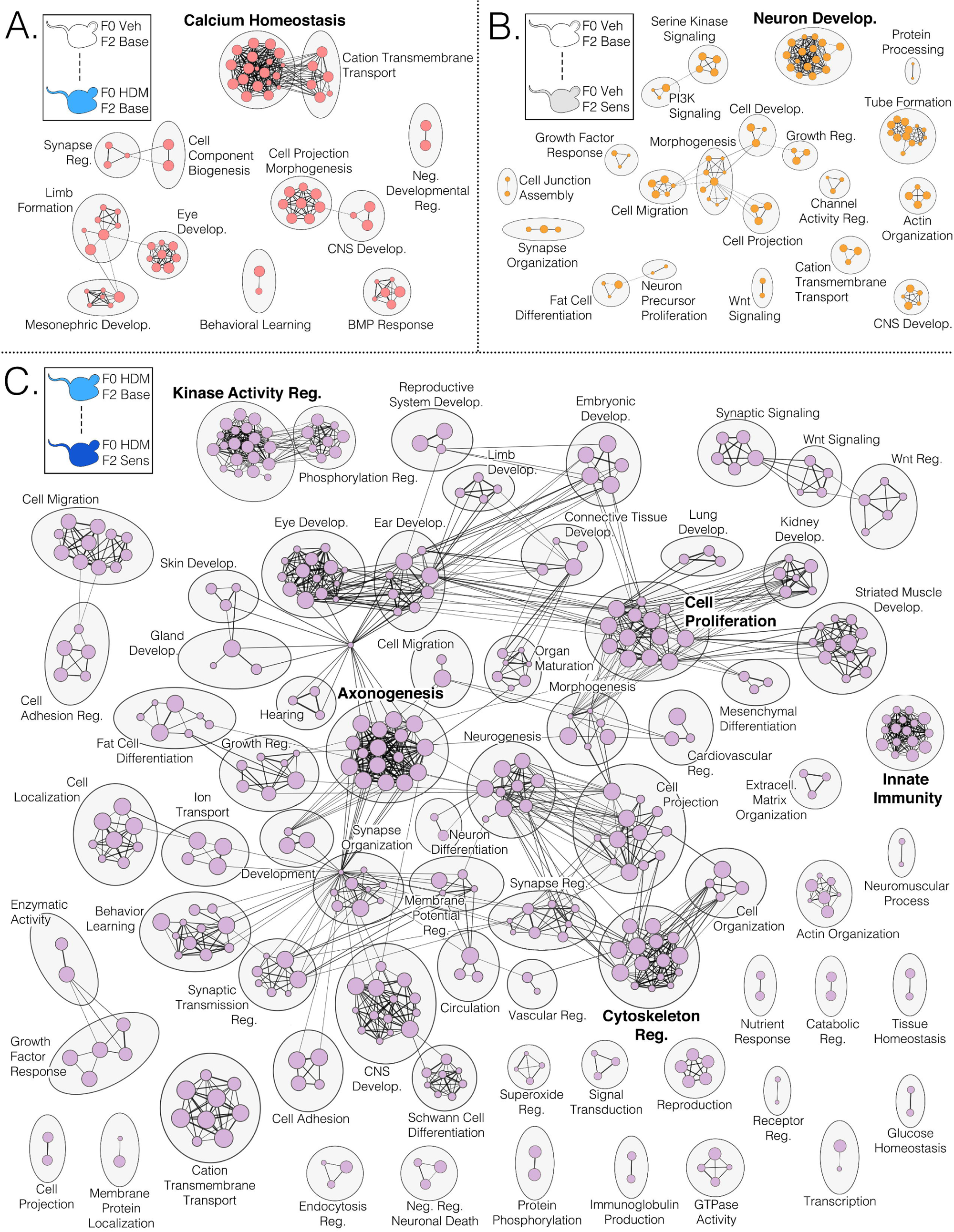
Gene ontology analysis of DMCs in airway epithelium of second-generation offspring. Gene ontology analysis of DMCs in airway epithelium of F2 mice identified enriched biological processes (individual small circles), which were then grouped based on functional annotation using Cytoscape Enrichment Map (larger, labeled, black circles). Each small circle represents an individual biological process and size is proportional to number of involved genes, while connecting edges represent shared genes between individual biological processes. (A) Gene ontology analysis showing enriched biological processes due to grandmaternal HDM exposure in F2 offspring at baseline (F0 HDM•F2 Base versus F0 VEH•F2 Base). (B) Gene ontology analysis showing effects of HDM sensitization alone on enriched biological processes in F2 offspring from vehicle-exposed F0 mice (F0 Veh•F2 Sens vs F0 VEH•F2 Base). (C) Gene ontology analysis showing potentiating effects of F0 grandmaternal HDM exposure on enriched biological processes after HDM sensitization in F2 mice (F0 HDM•F2 Sens vs F0 HDM•F2 Base).

**Table 1.**
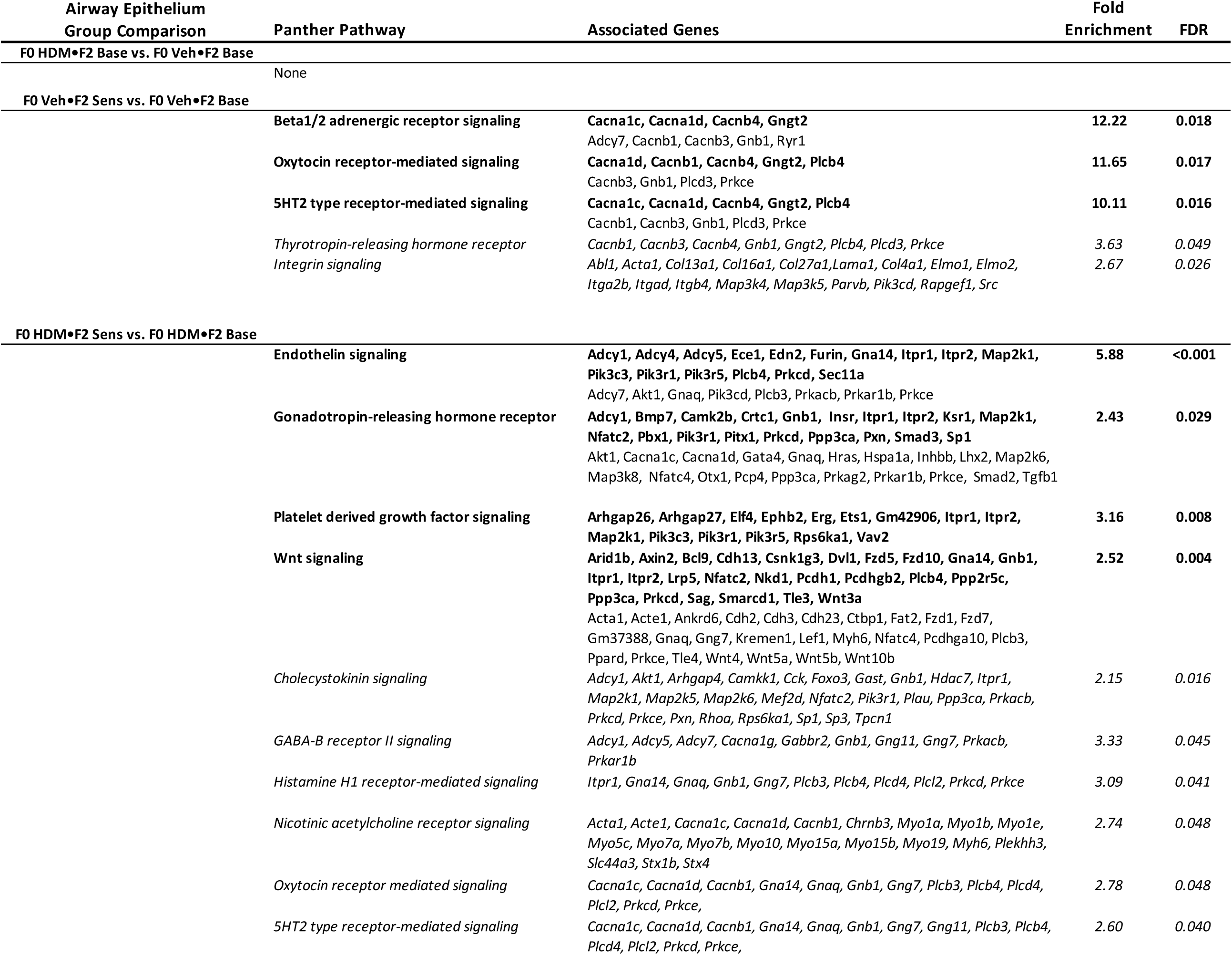
Pathway analysis of differentially methylated regions (DMR) and cytosines (DMC) in airway epithelium of second-generation mice. Pathways enriched in DMRs and genes associated with each pathway are denoted in bold font. All pathways identified in DMR analysis were similarly identified in DMC analysis. Additional pathways were identified in DMC analysis, denoted in italics, with associated genes.

### Allergen sensitization provokes a predominantly hypomethylated response in airway epithelium of second generation offspring from control founders

To test how allergen sensitization alone affected F2 methylation, DMRs and DMCs were assessed in airway epithelium of HDM-sensitized F2 mice (F0Veh•F2Sens) compared to non-sensitized F2 mice (F0Veh•F2Base), both from the vehicle-exposed F0 lineage. In total, 261 DMRs were identified that overlapped 277 unique genes. One hundred and sixty one (61.7%) were hypomethylated and 100 (38.3%) were hypermethylated. Most DMRs spanned intra- and intergenic sequences, with 20.3% mapping to promoters (**Figure 3**). We identified 1,362 DMCs, of which 754 (55.4%) were hypomethylated and 608 (44.6%) were hypermethylated. Similar to DMRs, most DMCs identified covered inter- and intragenic sequences, with 20.3% mapping to promoters (**Figure S1**).

Pathways associated with F0Veh•F2Sens DMRs included beta-1 and beta-2 adrenergic signaling, oxytocin receptor signaling, and 5HT2 serotonin receptor signaling pathways (**Table 1**). Additional pathways associated with DMCs included integrin signaling and thyrotropin-releasing hormone signaling pathways. Gene ontology analysis indicated that DMRs and DMCs were enriched for several biologic processes as well, including most notably, cell growth and migration, actin organization, and neuron development (**Figure 4B**).

### Grandmaternal allergen exposure potentiates DNA methylation and alters enriched pathways in allergen sensitized second generation offspring

To test whether grandmaternal allergen exposure alters F2 methylation after HDM sensitization, HDM sensitized F2 offspring (F0HDM•F2Sens) were compared to non-sensitized F2 offspring (F0HDM•F2Base), both from HDM-exposed F0 founders. A greater number of DMRs and DMCs were identified after allergen sensitization in F2 offspring from HDM-exposed founders compared to F2 offspring from vehicle-exposed founders. A total of 1,234 DMRs (Figure 3) and 3,756 DMCs (Figure S1) were identified in airway epithelium of F0HDM•F2Sens offspring, corresponding with 1,205 genes. Allergen sensitization also caused hypermethylation of most DMRs and DMCs in F0HDM•F2Sens offspring as compared to hypomethylation in F0Veh•F2Sens offspring. Of the 1,234 DMRs, 809 (65.6%) were hypermethylated, while 425 (34.4%) were hypomethylated. Most DMRs spanned introns, while 236 (19.1%) mapped to promoters (**Figure 3**) in F0HDM•F2Sens offspring. Of DMCs identified in this group, 2,261 (60.2%) were hypermethylated and 1,495 (39.8%) were hypomethylated. Similar to DMRs, most DMCs spanned introns, while 642 (17.1%) mapped to promoters (**Figure S1**). Pathway analysis of differentially methylated regions identified in F0HDM•F2Sens offspring included endothelin, platelet derived growth factor, Wnt, and gonadotropin-releasing hormone signaling pathways. Pathway analysis of DMCs also included enrichment for GABA-B receptor II, histamine H1 receptor, oxytocin receptor, nicotinic receptor, 5HT2 serotonin receptor, and cholecystokinin receptor signaling (**Table 1**).

Gene ontology analysis of DMCs demonstrated that grandmaternal allergen exposure changed enriched biological processes after allergen sensitization in F2 offspring. A total of 418 biological processes were enriched after allergen sensitization in F0HDM•F2Sens offspring compared with 97 after allergen sensitization in F0Veh•F2Sens (**Figure 4C**). After grouping functionally related GO terms, the most notably represented biological processes included axonogenesis, cell proliferation, innate immunity, cytoskeleton regulation and kinase activity regulation.

### Epigenetic changes in airway epithelial cells after allergen sensitization are uniquely regulated by grandmaternal allergen exposure

To test whether maternal HDM exposure affects DNA methylation after allergen sensitization in offspring, DMR-associated genes were compared in allergen sensitized F2 offspring from HDM-exposed and vehicle-exposed F0 founders (F0HDM•F2Sens vs. F0Base•F2Sens). In total, only 29 genes were in common between the 1,205 genes associated with DMRs in F0HDM•F2Sens mice and 277 genes associated with DMRs in F0Veh•F2Sens mice (**Figure 5A)**, indicating that maternal allergen exposure provokes a fundamentally unique methylation response after allergen sensitization in F2 mice. Furthermore, only 11 of 29 genes changed in the same direction (i.e. hypo- or hypermethylation), while the remaining 18 were differentially regulated between groups (**Figure 5B**).

**Figure 5.**
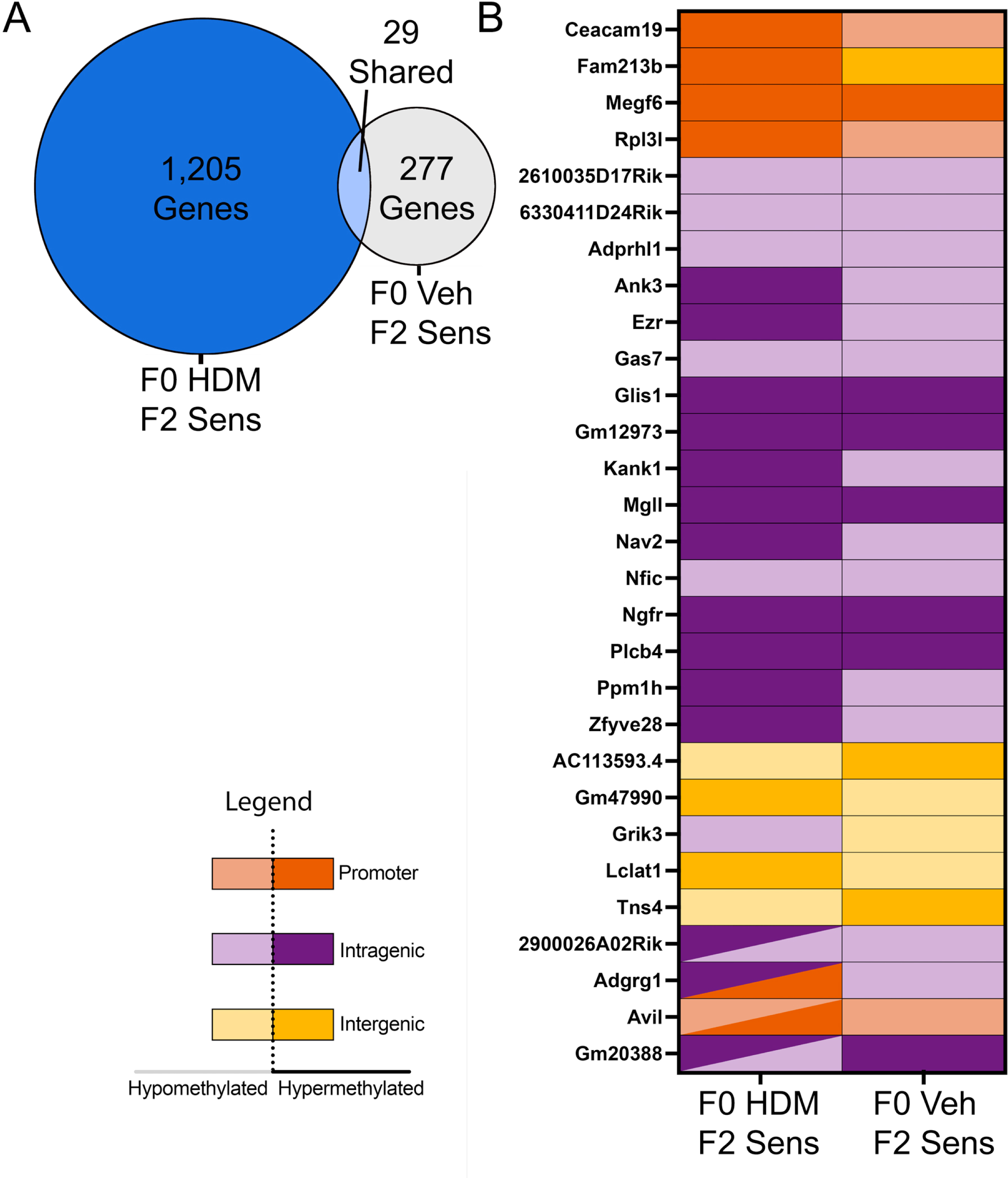
Unique genes are differentially methylated in airway epithelium after allergen sensitization in offspring from HDM-exposed founders. DMR-associated genes in HDM-sensitized F2 mice from HDM-exposed founders were compared against DMR-associated genes in HDM-sensitized F2 mice from vehicle-exposed founders to assess for common genes. (A) Twenty-nine genes were shared between F0 Veh•F2 Sens and F0 HDM•F2 Sens groups, out of 277 total genes in F0 Veh•F2 Sens mice and 1,205 total genes in F0 HDM•F2 Sens mice. (B) Of these 29 genes, 11 were similarly hyper- or hypomethylated between lineages, while 18 were differentially methylated.

### Multiple transcription factor binding sites are enriched in allergen-sensitized offspring from allergen-exposed founders

Since transcription factor binding sites regulate gene expression, enrichment of specific transcription factor binding sites within DMRs was analyzed in all groups. Several transcription factor binding sites were enriched in epithelial DMRs of allergen-sensitized offspring from allergen-exposed founders (F0HDM•F2Sens vs. F0HDM•F2Base) including several factors related to AP-1 factor family and p53 factor family (**Table 2**). In contrast, the few binding sites present in DMRs of allergen-sensitized F2 offspring from vehicle-exposed founders (F0Veh•F2Sens vs F0Veh•F2Base) and in non-sensitized F2 offspring from grandmaternal allergen exposed founders (F0HDM•F2Base vs F0Veh•F2Base) primarily related to the ETS family of factors.

**Table 2.**
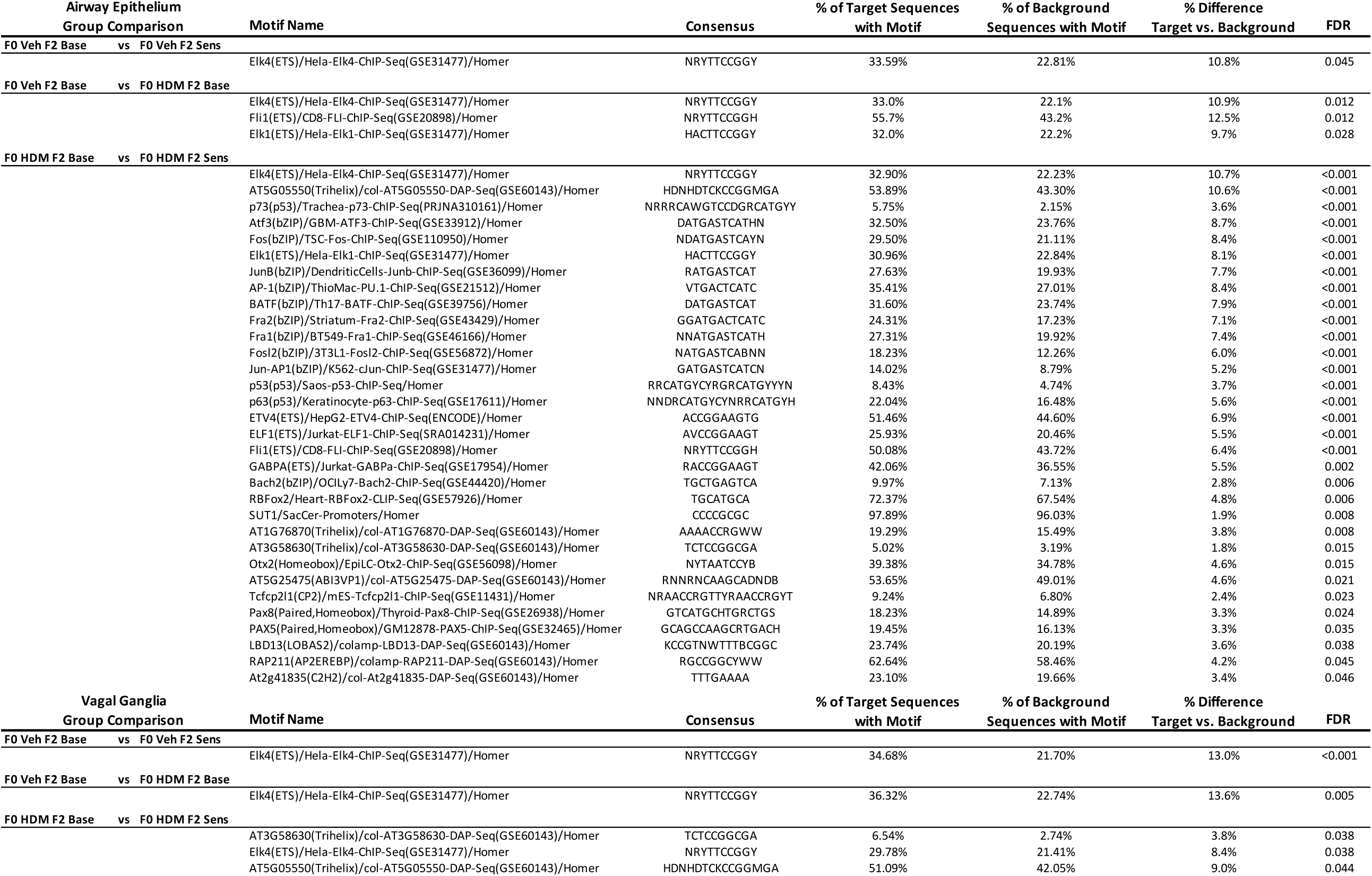
Over-represented transcription factor binding sites in DMRs of airway epithelium and sensory neurons from F2 mice before and after allergen sensitization.

### Grandmaternal allergen exposure influences epigenetic signatures of vagal ganglion sensory neurons

To test whether grandmaternal HDM exposure affected DNA methylation in vagal ganglia sensory neurons, F2 offspring from allergen-exposed F0 founders (F0HDM•F2Base) were compared to F2 mice from vehicle-exposed founders (F0Veh•F2Base). A total of 211 DMRs and 901 DMCs were detected in F0HDM•F2Base mice. Similar to airway epithelium, a majority of DMRs and DMCs in F0HDM•F2Base mice were hypermethylated, including 127 (60.2%) DMRs and 536 (59.5%) DMCs **(Figure 6 and Figure S2**). No pathways or biological processes were associated with differentially methylated regions or cytosines (DMRs or DMCs) in this comparison (**Table 3**).

**Figure 6.**
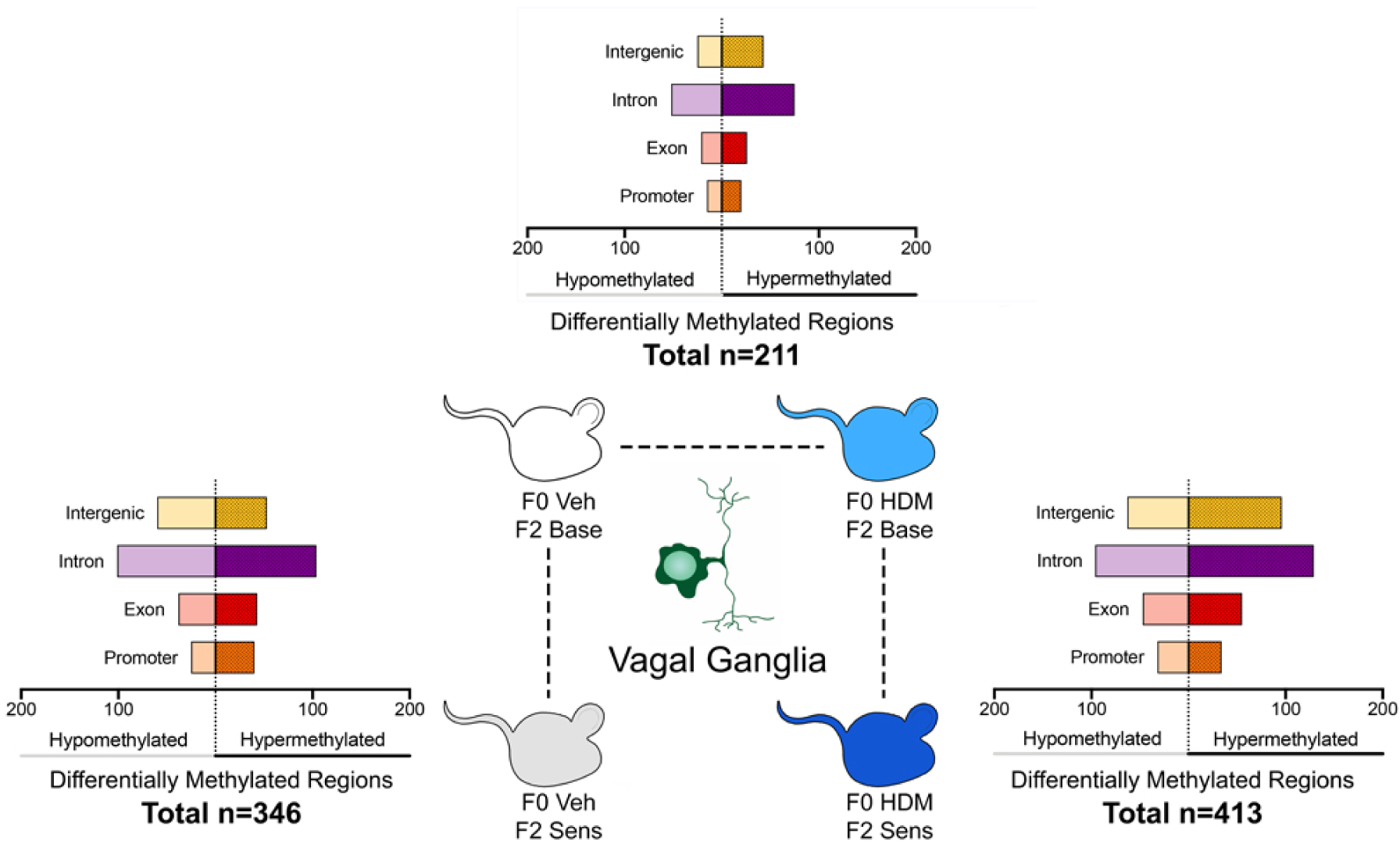
Grandmaternal allergen exposure alters vagal ganglion DNA methylation at baseline and after allergen sensitization in second-generation offspring. Grandmaternal HDM exposure alone resulted in 211 DMRs (F0 HDM•F2 Base versus F0 Veh•F2 Base) in vagal ganglia. F2 HDM sensitization alone resulted in 346 DMRs (F0Veh•F2Sens versus F0 Veh•F2 Base). In contrast, HDM sensitization of F2 mice from HDM-exposed founders resulted in 413 DMRs (F0 HDM•F2 Sens versus F0 HDM•F2 Base).

**Table 3.**
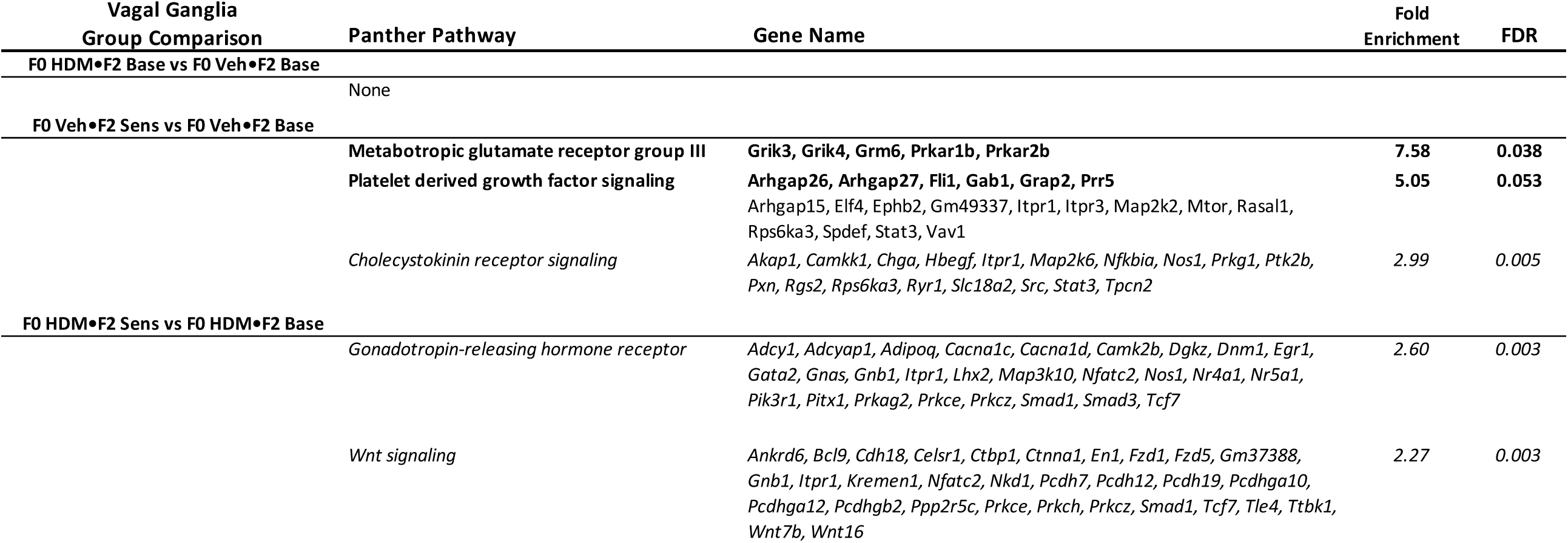
Pathway analysis of differentially methylated regions (DMR) and cytosines (DMC) in vagal ganglia of second-generation mice. Pathways enriched in DMRs and genes associated with each pathway are denoted in bold font. All pathways identified in DMR analysis were similarly identified in DMC analysis. Additional pathways were identified in DMC analysis, denoted in italics, with associated genes.

The 211 DMRs in vagal ganglia from F0HDM•F2Base mice corresponded to 222 unique genes, which were then compared with 297 genes detected in airway epithelium of the same mice. Only 14 genes were in common between vagal ganglia and airway epithelium, demonstrating that most epigenetic changes caused by grandmaternal HDM exposure occurred in a tissue-specific context.

Next, to determine the effects of allergen sensitization alone on vagal ganglia epigenetic signatures, HDM-sensitized F2 mice were compared to non-sensitized F2 mice from vehicle-exposed grandmothers (F0Veh•F2Sens vs. F0Veh•F2Base). A total of 346 DMRs and 1,467 DMCs were detected and were evenly balanced between hyper- and hypomethylation (**Figure 6 and Figure S2**), with 174 (50.3%) DMRs and 795 (54.2%) of DMCs being hypermethylated. Pathways associated with allergen sensitization in vagal ganglia of F0Veh•F2Sens included metabotropic glutamate receptor group III, platelet derived growth factor and cholecystokinin receptor signaling (**Table 3**). Gene ontology analysis indicated that several biologic processes were enriched, most notably axonogenesis, cell projection, cell adhesion and synapse organization (**Figure S3**).

Effects of grandmaternal HDM exposure on F2 methylation after HDM sensitization were determined by comparing HDM-sensitized F2 offspring from HDM-exposed founders (F0HDM•F2Sens) to non-sensitized F2 mice from HDM-exposed F0 founders (F0HDM•F2Base). Four hundred and thirteen DMRs and 1,787 DMCs were identified (**Figure 6 and Figure S2**), of which 242 (58.6%) DMRs and 892 (49.9%) DMCs were hypermethylated. No pathways were associated with DMRs, whereas, similar to findings in airway epithelial cells, gonadotropin-releasing hormone and Wnt signaling pathways were associated with DMCs (**Table 3**). Enriched biological processes included axonogenesis and transmembrane cation transport **(Figure S3)**.

### Epigenetic changes in vagal ganglia after allergen sensitization are altered by grandmaternal allergen exposure

DMR-associated genes were compared between allergen-sensitized F2 offspring from HDM-exposed and vehicle-exposed F0 founders (F0HDM•F2Sens vs. F0Veh•F2Sens). In total, 21 genes were shared between 222 genes associated with DMRs in F0Veh•F2Sens mice and 413 genes associated with DMRs in F0HDM•F2Sens mice (**Figure 7A)**. Of the 21 genes in common, only 7 had a matching pattern of hypo- or hypermethylation (**Figure 7B**).

**Figure 7.**
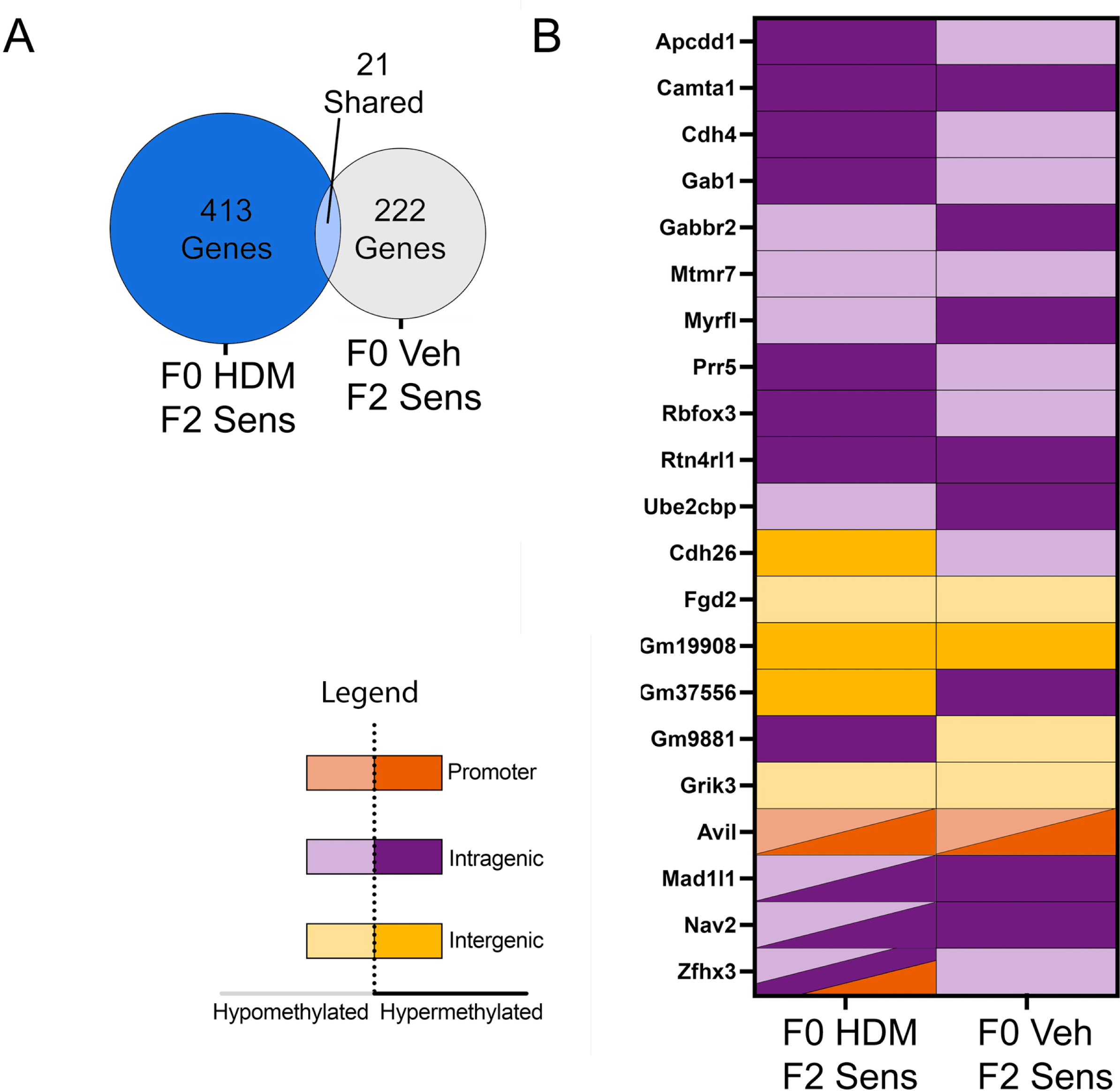
Unique genes are differentially methylated in vagal ganglia neurons after allergen sensitization in offspring of HDM-exposed founders. DMR-associated genes in vagal ganglia neurons were compared between HDM-sensitized F2 mice from HDM-exposed founders and HDM-sensitized F2 mice from vehicle-exposed founders (A) Twenty one genes were shared between F0 Veh•F2 Sens and F0 HDM•F2 Sens groups, out of 222 total genes in F0 Veh•F2 Sens mice and 413 total genes in F0 HDM•F2 Sens mice. (B) Of these 21 genes, only 7 had a matching pattern of hyper- or hypomethylated between lineages.

## DISCUSSION

In recent years, the epigenome has drawn intense interest as one potential explanation for the epidemic rise in asthma (DeVries and Vercelli, 2016). DNA methylation represents one epigenetic mechanism by which rapid phenotypic adaptations after environmental exposures can be transmitted from one generation to the next. Exposures to maternal factors during fetal lung development may be particularly impactful to asthma risk. However, linking specific maternal factors to epigenetic changes and developmental outcomes in humans has been challenging due to human longevity and confounding from a myriad of pre- and post-natal exposures. Maternal smoking has been most strongly linked with epigenetic changes associated with altered lung development and asthma risk (Hylkema and Blacquiere, 2009, Gilliland et al., 2000, Patil et al., 2013, Li et al., 2005). Maternal diet (Chatzi et al., 2008), stress (Gheorghe et al., 2010) and diesel exhaust exposure (Liu et al., 2008) have also been associated with alterations in the epigenome, and specific CpG methylation sites have been associated with atopy and asthma in children (Patil et al., 2013, Everson et al., 2015). Epigenetic changes regulate immune cell development including Th1, Th2, and T-regulatory cell differentiation (White et al., 2002, Fields et al., 2002, Baron et al., 2007) and differences in immune function can be detected at birth (Rinas et al., 1993, Ohshima et al., 2002), suggesting prenatal programming influences a child’s response to environment exposures from the earliest moments of life.

In animal models, particularly in mice, several factors are linked to transgenerational epigenetic regulation. For example, maternal diesel exhaust exposure altered dendritic cell DNA methylation that persisted across three generations and was associated with increased airway inflammatory responses to ovalbumin antigen in offspring (Gregory et al., 2017). A second study found that maternal HDM exposure altered methylation patterns in whole lung homogenates that were detectable in three successive generations and were associated with enhanced airway inflammation and airway responsiveness after HDM challenge (Pulczinski et al., 2021). Unlike this latter study, our analysis focused specifically on airway epithelium and vagal sensory neurons since methylation changes are highly cell specific (as shown by the different results we obtained in these two tissues). The central role of airway epithelium in the initiation of allergic asthma as both the site of contact for inhaled particles and as a key source of proinflammatory cytokines (Roan et al., 2019), and the important role of airway sensory neurons in regulating reflex bronchoconstriction and hyperreactivity make these two tissues of special interest.

Using the mouse model, we show that grandmaternal allergen exposure potentiates allergic airway inflammation and airway hyperreactivity in second-generation offspring. These effects were associated with significant alterations in offspring’s epigenetic signature at baseline and reprogramming of their epigenetic response to allergen sensitization later in life, in a tissue-specific context.

In our experiments, DMRs and DMCs associated with allergen sensitization alone (i.e. allergen-sensitized F2 mice from the control lineage) mapped to voltage-gated L-type calcium channels involved in multiple G protein coupled receptor signaling pathways (i.e. beta adrenergic receptors, 5-HT2 receptors, oxytocin receptors and thyrotropin-releasing hormone receptors), and to the integrin signaling pathway, both of which contribute to normal airway function. L-type calcium channels on epithelium regulate mucin secretion (Khakzad et al., 2012) and are involved in branching morphogenesis during development (Brennan et al., 2013), whereas airway integrins are essential for cellular adhesion and maintenance of epithelial barrier function (Wright et al., 2014). Loss of epithelial barrier integrity has been implicated in asthma pathogenesis and specific integrins that were differentially methylated in our study, such as integrin B4, are down-regulated in humans with asthma (Wu et al., 2020, Liu et al., 2010). Mice with integrin B4 deficiency exhibit accelerated epithelial cell senescence (Yuan et al., 2019) and have an exaggerated inflammatory response to allergen (Liu et al., 2018). Inhibition of other integrin signaling mediators identified in our analysis, such as PiK3, also result in potentiated inflammatory responses and airway hyperreactivity in mice (Lee et al., 2006), suggesting that dysregulation at multiple levels of integrin signaling may exacerbate responses to allergens.

Importantly, our data also show that grandmaternal allergen exposure fundamentally alters future generations epigenetic response to allergen sensitization. In allergen-sensitized offspring from allergen-exposed founders, DMRs and DMCs mapped to several signaling pathways that regulate lung development and inflammation, including WNT signaling, platelet derived growth factor signaling, gonadotropin-releasing hormone receptor pathway, and endothelin receptor signaling. Differentially methylated WNT proteins in our model included LRP5 and Axin2, which regulate airway branching and alveolar morphogenesis in developing fetal lungs (Mammoto et al., 2012, Frank et al., 2016, Zhang et al., 2012). Other differentially methylated genes coding WNT proteins, such as Nfatc2 (Jakobi et al., 2020, Hinds et al., 2013) and PCDH1 (Koppelman et al., 2009) have been identified as novel susceptibility loci for development of allergy and bronchial hyperreactivity in humans, respectively. *PCDH1* polymorphisms were associated with development of asthma in children (Mortensen et al., 2014, Faura Tellez et al., 2015) and *PCDH1* function was critical to epithelial barrier function in vitro (Kozu et al., 2015). Expression of *WNT3, WNT5a, WNT10* and their receptor, Fzd-5, positively correlated with type 2 high asthma in adults while expression of WNT5b was negatively correlated with a type 2 signature (Choy et al., 2011). Tle3 regulated T cell development (Xing et al., 2018) and loss of WNT10b enhanced airway inflammation in a murine model (Trischler et al., 2016). Functionally, WNT signaling is critical to epithelial–mesenchymal differentiation during development (Sharma et al., 2010, Dean et al., 2005) and at times of lung injury (Konigshoff and Eickelberg, 2010), where their ability to broadly influence transcriptional events regulates airway inflammation and epithelial repair.

Other over-represented pathways in allergen-sensitized offspring of allergen-exposed founders included PDGFR signaling and endothelin receptor signaling. PDGF family proteins are involved in cell proliferation and migration during embryogenesis and contribute to airway remodeling by inducing epithelial metaplasia and airway smooth muscle migration in later life (Cohen et al., 2009). PDGF mediators MEK1 and Pi3k contributed to airway epithelial cell differentiation during development (Boucherat et al., 2014) and mediated epithelial cytokine production after HDM exposure in adult mice (Kim et al., 2020b). Consequently, blocking PI3K reduced aspergillus-induced allergic airway inflammation in mice (Jeong et al., 2018). Similarly, endothelin pathway mediators played an important role initiating eosinophilic inflammation after allergen (Finsnes et al., 1998). In rats, endothelin levels rapidly increased in bronchoalveolar lavage fluid and in airway epithelium after allergen exposure and airway inflammation was blocked by endothelin receptor antagonists (Finsnes et al., 1997). The endothelin signaling protein ITPR2 is important for cellular adhesion as well as T and B cell activation (Patterson et al., 2004) and ITPR2 polymorphisms are associated with asthma in humans (Wilker et al., 2009). Endothelin also has the ability to directly induce human airway smooth muscle contraction (Advenier et al., 1990). Thus, differential methylation in our study has identified pathways involving airway immune responses, lung development, and airway contractility.

Gene ontology analysis demonstrated greater enrichment of biological processes in allergen-sensitized offspring from the allergen-exposed lineage. Foremost among these processes were cell proliferation, regulation of kinase activity, innate immunity, axonogenesis and cytoskeleton regulation, all of which were parts of a larger theme of cellular migration and organ morphogenesis. In contrast, allergen sensitization alone (i.e. allergen-sensitized offspring from control founders), and effects of maternal allergen exposure alone (i.e. control offspring from allergen-exposed F0 mice), affected only a small fraction of developmental processes.

Several transcription factor binding sites were enriched in epithelial DMRs of allergen-sensitized offspring from allergen-exposed founders, including members of the AP-1 and p53 factor families. Transcription factor binding sites, which are typically located near promoters, increase or decrease gene transcription. It is therefore possible that these DMRs act as regulatory sites and their differential methylation might impact accessibility to binding from transcription factors (Tulchinsky et al., 1996). AP-1 consists of Fos (c-Fos, FosB, FosL1, FosL2) and Jun (c-Jun, JunB, JunD) proteins, which form dimers that regulate transcription of inflammatory genes relevant to asthma, such as c-Fos and interleukin (IL)-5, and Fra2 and IL-13 (Liu et al., 2004, Gungl et al., 2018, Ferreira et al., 2020). AP-1 binding also reduces glucocorticoid receptor binding site availability by altering chromatin structure (Adcock et al., 1995, Jacques et al., 2010), suggesting AP-1 has a role in development of corticosteroid-resistant disease. Accordingly, in asthma and particularly in corticosteroid-resistant asthma, expression of AP-1 dimers (e.g. c-Fos) is increased (Sousa et al., 1999, Lane et al., 1998). AP-1 can also be activated by inflammatory cytokines TNF-alpha and IL-1beta, suggesting transcription factor activity is amplified by its own downstream products.

The p53 transcription factor family includes structurally similar factors p53, p63, and p73. While more commonly known for their role as tumor suppressors, p53 and p63 are involved in epithelial cell senescence and epithelial wound repair, respectively (Warner et al., 2013, Moheimani et al., 2015). Both of these processes are implicated in asthma pathogenesis. p73 also has a role in mucociliary development. Thus, p53-associated factors have several potential implications for lung development and disease pathogenesis.

Our analysis included an evaluation of methylation in airway sensory neurons since 1) airway nerves control bronchoconstriction (Drake et al., 2021) 2) sensory nerve remodeling correlated with worse lung function and increased irritant sensitivity in humans with eosinophilic asthma (Drake et al., 2018) and 3) maternal type-2 inflammation in mice increased airway sensory nerve density and nerve-mediated airway hyperreactivity after allergen exposure in first generation offspring (Lebold et al., 2019). Grandmaternal allergen exposure significantly altered methylation in sensory nerves at baseline and after allergen sensitization. In total, only 21 genes corresponding to DMRs were shared between allergen-sensitized offspring from allergen-exposed versus control lineages, lending further evidence that grandmaternal allergen exposure results in a substantial tissue specific reprograming of epigenetic responses to allergen. While individual genes associated with DMRs varied markedly between vagal ganglia and airway epithelium in F2 offspring from allergen-exposed founders, enriched pathways were similar, including PDGF signaling, gonadotropin releasing hormone receptor signaling, and WNT signaling pathways. Additional involvement of cholecystokinin receptor signaling in vagal ganglion at baseline occurred as the result of grandmaternal allergen exposure. Cholecystokinin is a typically associated with sensory nerve activation in the gut, with less known about its role in sensory neurons in airways. Cholecystokinin expression in neuroendocrine bodies has been detected in lower airways in sheep (Balaguer et al., 1992) and airway sensory nerves increase cholecystokinin receptor expression after lung injury (Kaelberer et al., 2020), possibly resulting in increased bronchoconstriction in humans (Stretton and Barnes, 1989).

In summary, our study indicates that grandmaternal allergen exposure increases airway reactivity and inflammation in offspring after allergen sensitization and challenge, in association with reprogramming of epigenetic responses to allergen sensitization. We showed that epigenetic trajectories are established before birth in response to prenatal exposures, which direct subsequent epigenetic responses to allergen sensitization and contribute to transgenerational inheritance of asthma risk.

## METHODS

### Mice

Male and female C57Bl/6J mice (Jackson Laboratories, Bar Harbor, ME) were 8-14 weeks old at time of experimentation. Mice were housed in filtered air rooms with ad libitum access to food and water on a 12-h light/dark cycle and were treated in accordance with standards established by the United States Animal Welfare Act set forth by the National Institutes of Health guidelines. The Institutional Animal Care and Use Committee at Oregon Health & Science University approved all experimental protocols.

### Grandmaternal (F0) Allergen Exposure

Nulliparous female mice were exposed to 25 μg house dust mite (HDM, dissolved in 25 μL PBS, Greer Laboratories, n=10 mothers) or vehicle (PBS, n=10 mothers) for 7-8 total weeks. For the first 4 weeks, mice were sedated with 5% isoflurane and intranasally exposed to HDM or vehicle once daily on 5 consecutive days per week (Figure 1). At the end of week 4, females were paired with untreated males for breeding. From week 5 until delivery, pregnant females were manually restrained without isoflurane sedation, since pilot studies showed daily sedation caused fetal demise, and exposed intranasally with 25 μg HDM or vehicle, 5 days per week through gestation until birth. Maternal treatments ceased at time of delivery.

### First Generation Offspring (F1) Breeding Scheme

First generation offspring (F1) were randomly mated with offspring born to different F0 mothers from the same treatment group (HDM or Veh) at 8 weeks of age. Thus, each mated pair represented an F1 male offspring and an F1 female offspring from different F0 mothers, resulting in n=5 mating pairs. F1 mice were not sensitized with HDM and received no other treatments.

### Second Generation Offspring (F2) HDM Sensitization and Challenge

F2 offspring mice 8-12 weeks of age were anesthetized with 5% isoflurane and sensitized with 50 μg HDM i.n. or vehicle on days 0 and 1. On day 14, one cohort of mice were utilized for methylation analysis while a second cohort underwent allergen challenge with 25 μg HDM or vehicle i.n. on days 14-17. Airway physiology and inflammation were measured on day 18.

### Assessments of Airway Physiology and Inflammation

F2 mice airway physiology was measured as previously described (Lebold et al., 2019). Briefly, mice that were randomly assigned for physiologic studies were sedated with ketamine (100 mg/kg i.p.) and xylazine (10 mg/kg i.p.), paralyzed with succinylcholine (10 mg/kg i.p.), tracheotomized, and mechanically ventilated via a 21-gauge catheter. A constant volume ventilator delivered 200 µL tidal volumes at 125 breaths/min with an inspired oxygen content of 95% oxygen/5% CO2 and 2 cm H_2_O of positive end-expiratory pressure. Flow and pressure were continuously monitored (AD Instruments) and aerosolized serotonin (10-1000 mM; AeroNeb, Torrington, CT) was delivered at 2 minute intervals. Airway resistance, calculated as the difference between peak inspiratory pressure and plateau pressure during an end inspiratory pause, divided by airflow (Resistance = Ppeak-Pplateau / flow) was measured 60 seconds after each dose of serotonin and expressed as fold change over resistance to aerosolized PBS.

Bronchoalveolar lavage was performed by instilling 0.5 ml sterile PBS three times into the lungs via the tracheal cannula. Total cells were determined with a hemacytometer and cell differentials measured using Wright staining.

### DNA Isolation, Extraction and Purification

Mice that were randomly assigned for DNA isolation were injected with a lethal dose of pentobarbital (150 mg/kg i.p.) and perfused with sterile PBS. Tracheas and primary bronchi were excised and incubated in 0.5% protease solution in RPMI medium containing penicillin-streptomycin-amphotericin B at 4°C overnight, followed by fresh protease solution at 37°C for 30 minutes. Airway epithelium was then isolated by instilling sterile PBS through the tracheal lumen. Isolated cells were counted using a hemocytometer and >95% cell viability was confirmed with Trypan Blue. Concurrently, mouse vagal ganglia were removed and placed in sterile cell lysis buffer (Qiagen Puregene, Hilden Germany) with 0.5% SDS and 100ug/ml Proteinase K at 55°C overnight. Both epithelium and vagal sensory neurons were stored in sterile cell lysis buffer prior to DNA extraction.

DNA were extracted from airway epithelial cells and vagal sensory neurons using Gentra Puregene Tissue (Qiagen) and purified using gDNA Clean and Concentrator 10 columns (Zymo Research, Irvine, CA) by the Gene Profiling Shared Research Core at OHSU. DNA concentration and purity were assessed by UV absorption (Nanodrop One, ThermoFisher, Waltham, MA), size analysis was performed using the gDNA Tape method (2200 TapeStation, Agilent, Santa Clara, CA), and quantification was performed using PicoGreen fluorescence assay (Sigma, Burlington, MA).

### Reduced-representation bisulfite sequencing (RBSS) library generation

RRBS libraries were generated as previously described (Carbone et al., 2019). Briefly, DNA (100-150 ng) from epithelium or sensory neurons underwent digestion for 10 hours at 37^°^C with *MspI* restriction enzyme (New England Biolabs, Ipswich, MA), followed by purification with AMPure XP (Beckman Coulter, Indianapolis, IN). Libraries were generated with a NEXTflex Bisulfite-Seq Kit (Bioo Scientific Corporation, Austin, TX) paired with the NEBNext Methylated Adaptor (New England Biolabs). Bisulfite conversion was performed with a EZ DNA Methylation-Gold Kit (Zymo Research) followed by PCR amplification with NEBNext Multiplex Oligos (New England Biolabs). Libraries were quantified with the Qubit High Sensitivity dsDNA Assay (Life Technologies, Eugene, OR), and multiplexed for sequencing on the Illumina NextSeq 500 with the high-output, 75-bp cycle protocol.

### Differential methylation analysis

RRBS reads were analyzed for quality with FastQC (v0.11.5), followed by trimming with TrimGalore (v0.5.0) with the “—rrbs” parameter specified. Trimmed reads were aligned to the Ensembl mouse reference genome (GRCm38) with Bismark (v0.19.0) (Krueger and Andrews, 2011) using default parameters. Alignment rates were approximately 75%, consistent with typical bisulfite-converted sample alignment. For differentially methylated cytosines (DMC) analysis, only CpGs with at least 10X coverage and less than the 99.9^th^ percentile of the highest coverage CpGs in at least 4 replicates per group in a given tissue were considered. In total, 514,227 CpGs were used for epithelial DMC analysis and 541,043 CpGs were used in vagal sensory neuron DMC analysis. DMC were identified using a logistic regression model that utilized a Chi-square test, taking biological replicates into account and considering sex as a covariate in the model to calculate P-values. P-values were corrected to q-values using the SLIM method. A q value < 0.1 and an absolute methylation percent difference > 10% was considered significant. Differentially methylated region (DMR) analysis was performed by comparing non-overlapping 1000bp segments between experimental and reference genomes using logistic regression, as previously described (Akalin et al., 2012). Genes associated with DMCs and DMRs were annotated using Ensembl annotation GRCm38.92 and a custom script that utilized BEDTools (Quinlan and Hall, 2010) and the genomation R library (Akalin et al., 2015). For intergenic DMRs, the closest gene and the distance between the DMC/DMR and Transcription Start Site (TSS) was also annotated.

### Gene Ontology and Pathway Analysis

Gene ontology (release date 20210101) and pathway analysis of significant DMCs and DMRs were performed with Panther (release date 20200728). Hypo- and hypermethylated regions were jointly analyzed using overrepresentation with Fischer’s exact test and FDR correction (FDR<0.05). Over and underrepresented biological process GO terms with >5 and <1000 genes were visualized using Cytoscape Enrichment Map and shared gene clusters with functionally related GO terms were labeled.

### Transcription Factor Binding Site Analysis

Enrichment of transcription factor binding sites within significant DMRs was performed using HOMER Motif Enrichment (Heinz et al., 2010), specifically the “findMotifsGenome” script. The percentage of target sequences with a given motif were compared to the percentage of background sequences with a given motif. Only sequences with a q-value < 0.05 were considered significant.

### Statistics

Airway physiology data were analyzed with a repeated-measures two-way ANOVA with a Tukey’s post-test using Prism (Graphpad). Physiologic data that were greater than 2 standard deviations above or below their gender and group mean were considered outliers and excluded from analysis. Final N for each group were as follows: F0Veh•F2Veh n=13, F0Veh•F2HDM n=20, F0HDM•F2Veh n=13, F0HDM•F2HDM n=18. Bronchoalveolar lavage cell count data were analyzed with a two-way ANOVA and Tukey’s post-test. Hypo- and hypermethylated regions were jointly analyzed using overrepresentation with Fischer’s exact test and FDR correction (FDR<0.05). Genomics data was assessed using R as described above.

## SUPPLEMENTARY FIGURE LEGEND

**Supplementary Figure 1. Grandmaternal allergen exposure regulates differentially methylated cytosines in airway epithelium of second-generation offspring at baseline and after allergen sensitization**. Maternal HDM exposure alone resulted in 1,390 DMCs in airway epithelium of F2 offspring (F0 HDM•F2 Base vs F0 Veh•F2 Base). F2 HDM sensitization alone resulted in 1,467 DMCs (F0Veh•F2Sens vs F0 Veh•F2 Base). In contrast, HDM sensitization in F2 mice from HDM-exposed founders resulted in 3,756 total DMCs (F0 HDM•F2 Sens vs F0 HDM•F2 Base).

**Supplementary Figure 2. Grandmaternal allergen exposure regulates vagal ganglia cytosine methylation in second-generation offspring at baseline and after allergen sensitization**. Grandmaternal HDM exposure alone resulted in 901 DMCs (F0 HDM•F2 Base vs F0 Veh•F2 Base). F2 HDM sensitization alone resulted in 1,467 DMCs (F0Veh•F2Sens vs F0 Veh•F2 Base). In contrast, HDM sensitization in F2 mice from HDM-exposed maternal founders resulted in 1,787 total DMCs (F0 HDM•F2 Sens vs F0 HDM•F2 Base).

**Supplementary Figure 3. Gene ontology analysis of biological process enrichment in vagal ganglia from second-generation offspring**.

Gene ontology analysis of DMCs in vagal ganglia of F2 mice identified enriched biological processes (individual small circles), which were then grouped based on functional annotation using Cytoscape Enrichment Map (larger, labeled, black circles). Each small circle represents an individual biological process and size is proportional to number of involved genes, while connecting edges represent shared genes between individual biological processes. (A) Gene ontology analysis showing effects of HDM sensitization on biological process enrichment in F2 offspring from vehicle-exposed F0 mice (F0 Veh•F2 Sens vs F0 VEH•F2 Base). (B) Gene ontology analysis showing potentiating effects of F0 grandmaternal HDM exposure on biological process enrichment in F2 mice after HDM sensitization (F0 HDM•F2 Sens vs F0 HDM•F2 Base).

## Notes

### Competing Interest Statement

The authors have declared no competing interest.

