## Supplementary figures and images for "Grandmaternal allergen exposure causes distinct epigenetic trajectories in offspring associated with airway hyperreactivity and inflammation"

### Supplemental Figure 1

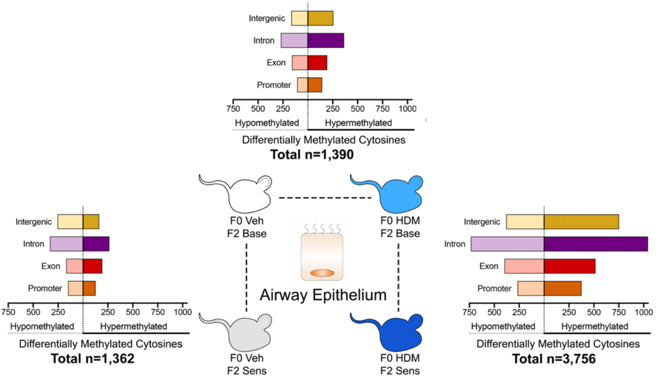

### Supplemental Figure 2

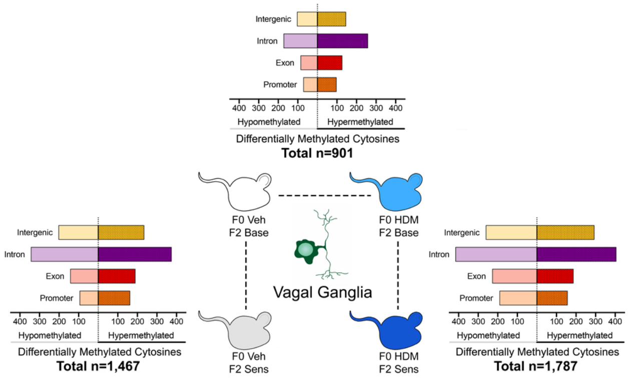

### Supplemental Figure 3

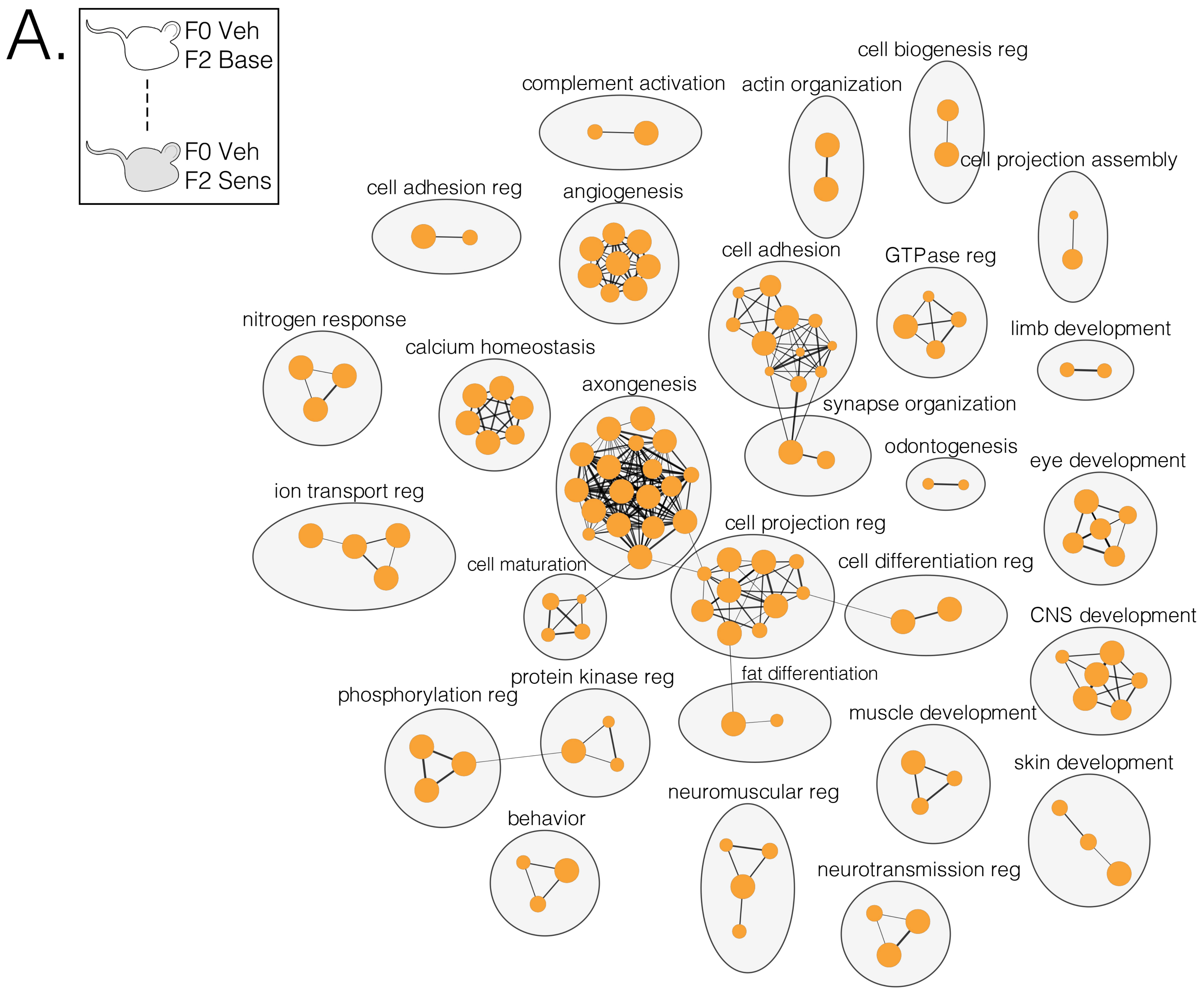

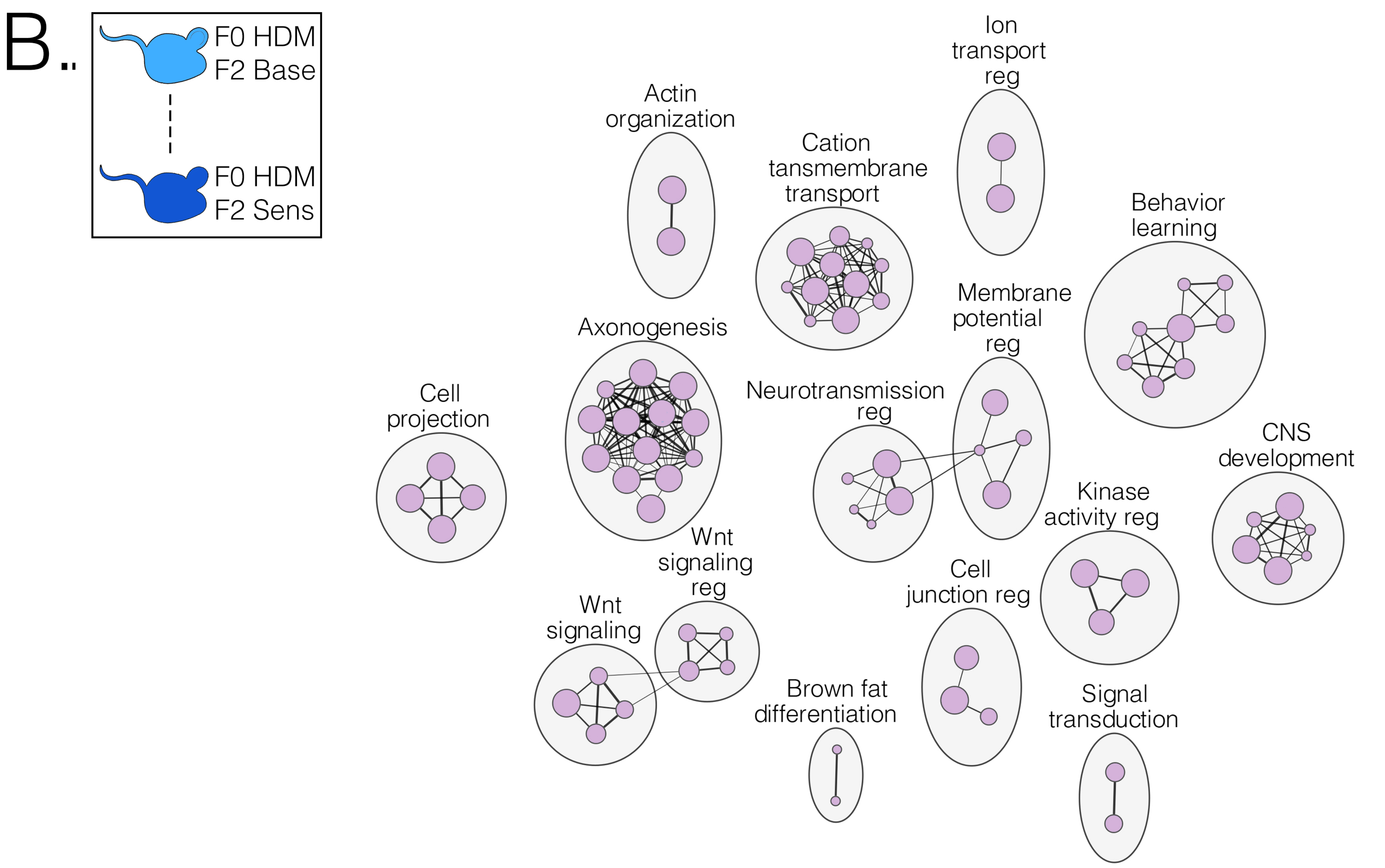
